# Rethinking ratio-based normalization: A guide towards model-based approaches in heart weight analysis

**DOI:** 10.1101/2025.09.01.673426

**Authors:** Manuela A. Oestereicher, Patricia da Silva-Buttkus, Valérie Gailus-Durner, Susan Marschall, Helmut Fuchs, IMPC consortium, Martin Hrabě de Angelis, Elida Schneltzer, Nadine Spielmann

## Abstract

Heart weight is a critical parameter in cardiology and mouse research, reflecting structural and functional changes linked to cardiac size or hypertrophy and pathophysiological conditions. Normalizing heart weight (HW) to body weight (BW) or tibia length (TL) is a common practice; however, the validity of these ratios has been questioned due to non-proportional relationships between parameters, and this becomes particularly problematic when comparing distinct populations based on such normalized values. Using data from over 25,000 C57BL/6N wildtype mice provided by the International Mouse Phenotyping Consortium (IMPC), we investigated the limitations of ratio-based normalization when comparing different groups, aiming to propose a robust framework for HW analysis.

Our findings reveal negligible to weak correlations between HW, BW, and TL across age and sex groups, undermining the validity of ratio-based methods. A modelling study using simulated data demonstrated that ratios could produce misleading results, including reversed or false group differences, when scaling assumptions are violated. Ratios yield accurate and interpretable results only when a truly proportional relationship exists between the variables—specifically, when the regression line passes through the origin—conditions under which ratio-based normalization aligns with outcomes obtained from more robust modelling approaches.

These results underscore the superiority of linear models with covariate adjustment and allometric scaling for organ weight analysis, as they more accurately capture biologically relevant scaling relationships. By leveraging the IMPC’s large-scale wildtype dataset, we establish the necessity of reassessing normalization practices in quantitative biology traits and propose that ratios should be avoided when comparing normalized values across distinct populations unless key mathematical assumptions are met. This study advances the analytical rigor in phenotyping research, enabling more accurate interpretations of organ mass and function across biological contexts.

## Introduction

Heart weight (HW) is a fundamental parameter in both human cardiac pathology and mouse research of the heart, providing insights into structural and functional changes in the heart. In clinical cardiology, increased HW, often indicative of hypertrophy, serves as a key marker of pathophysiological conditions such as hypertension and heart failure, reflecting adaptations that may progress to disease^1^.

Accurate assessment of the heart is essential for distinguishing pathological changes from normal variants. However, overlapping normal and abnormal ranges, often influenced by body size, pose challenges for clinical decisions, prompting a reassessment of normalization practices in cardiovascular research^2,3^.

In preclinical research, particularly in mice, HW is routinely measured to evaluate the effects of genetic, pharmacological, or environmental factors on cardiac health. The heart weight-to-body weight (HW/BW) ratio is the most commonly used normalization method, aiming to account for body size differences^4^. Normalization to tibia length (HW/TL) has also gained traction, as it better reflects skeletal growth and is less influenced by acute body weight fluctuations^5^. These ratios are widely used to discern whether heart weight variations are driven by pathological changes in heart weight or body size^6^.

Recent studies have raised concerns about the mathematical validity of using these ratios to compare populations^7^, particularly with non-proportional relationships between HW and body size metrics^8,9,10^. This has prompted efforts to develop alternative methods that prevent pseudo-indexing in animal models^11^. To address these issues, we analysed data from over 25,000 C57BL/6N wildtype control mice phenotyped by the International Mouse Phenotyping Consortium (mousephenotype.org). This large-scale dataset enabled us to systematically evaluate the limitations of ratio-based normalization and propose a robust statistical framework for analysing HW. Allometric scaling offers a robust framework for normalizing heart weight relative to body dimensions in mice, addressing the inherent non-linear relationships between the two. This method requires key steps, including determining the proportionality factor and allometric exponent. These parameters define the scaling relationship and ensure accurate normalization across experimental groups. To refine this approach, we applied linear models with covariate adjustments and allometric scaling, which were stratified by sex and age.

Our analysis demonstrates that linear and allometric models, depending on data structure, more effectively capture scaling relationships between biological traits and organism size, and offer a more robust framework for normalization and comparative studies.

## Methods

### The International Mouse Phenotyping Consortium

The International Mouse Phenotyping Consortium (IMPC) represents a multi-institutional and collaborative research initiative encompassing twenty-four major research organizations and funding agencies, distributed globally^12,13^. The IMPC seeks to generate and phenotype a knockout mouse line for every protein-coding gene with a human ortholog in the mouse genome (www.mousephenotype.org)^14^. Phenotyping is carried out under the standard operating procedures detailed in IMPReSS (International Mouse Phenotyping Resource of Standardized Screens: www.mousephenotype.org/impress/index), developed and validated during the pilot programs EUMORPHIA and EUMODIC^15^.

### IMPC centres contributing heart weight, body weight and tibia length data

IMPC data release (DR) 21.0 was used herein (https://www.mousephenotype.org/data/release). The following subset of eight IMPC data-contributing centres provided heart weight (HW) and body weight (BW) whereas tibia length (TL) data was provided by three centres (HMGU, ICS and KMPC) early adult (EA) and late adult (LA) and BCM for EA only in DR 21.0 (ethical approval details are included in parenthesis following the name of each contributing centre):

1. Baylor College of Medicine (BCM) (Institutional Animal Care and Use Committee approved license AN-5896).
2. German Mouse Clinic, Helmholtz Zentrum München (GMC) (#144-10, 15-168)
3. The Jackson Laboratory (JAX) Institutional Animal Care and Use Committee approved licenses 14004, 11005, and 99066. JAX AAALAC accreditation number was 000096, NIH Office of Laboratory Animal Welfare assurance number was D16-00170).
4. Institute Clinique de la Souris, Mouse Clinical Institute (ICS) (#4789-2016040511578546v2).
5. RIKEN BioResource Research Center (RBRC) (Animal Care Committee approved animal use protocols 0153, 0275, 0277, and 0279).
6. University of California – Davis (UCD) (Institutional Animal Care and Use Committee approved animal care and use protocol number 19075. UCD AAALAC accreditation number is 000029, and the NIH Office of Laboratory Animal Welfare assurance number is D16-00272 # (A3433-01).
7. Seoul National University, Korea Mouse Phenotyping Center (KMPC) (KRIBB-AEC-19189).
8. The Centre for Phenogenomics, Toronto (TCP) (0275 and 0279).

### Animals

This study includes data collected from inbred wildtype control animals tested as part of the IMPC goals. These mice, both males and females, were on a C57BL/6N genetic background of substrains: C57BL/6NCrl (HMGU, ICS and TCP); C57BL/6NJ (BCM, JAX), C57BL/6NJcl (RBRC) and C57BL/6NTac (HMGU, ICS, KMPC and RBRC).

HW, BW, and TL data were collected from mice at one of two possible time points. For the EA pipeline data were collected at a median age of 16 weeks with a minimum of 15 and maximum of 20 weeks of age. For the LA pipeline data were collected at a median age of 60 weeks with a minimum of 55 and maximum of 81 weeks of age. Animal welfare was assessed routinely during the phenotyping pipeline for all mice.

### Heart and body weight measurements

The IMPC standard operating procedure provides an overview of the heart and body weight procedures used by contributing centres:

https://www.mousephenotype.org/impress/ProcedureInfo?action=list&procID=601.

In brief, body weight (g) was recorded and then the mouse humanely euthanized. The mouse was placed in the supine position and exposed fur was wiped with 70% ethanol to control dander. Centre-specific complete necropsy and tissue collection according to technical SOPs were performed. This included removal of the heart by dissecting the aortic root immediately above the aortic valves and the superior vena cava above the atria. Carefully removing adjacent mediastinal fat pads from the excised heart with forceps and emptying the heart of blood by repeatedly tapping the heart on a Kimwipe (absorbent pad) or surgical compress until the heart was totally empty of blood. Heart weight (in mg) was recorded in the centre-specific database prior to immersing the heart in a fixative solution. All data were collected at a local workstation in the necropsy room (attached to a digital balance) and uploaded to the centre-specific pathology data capture system. Data were then uploaded to the IMPC data coordination centre for quality control, processing and analysis prior being released on the IMPC portal.

### Measurement of mouse tibia length

The IMPC standard operating procedure provides an overview of the tibia lengths (TL) procedure used by contributing centres:

https://www.mousephenotype.org/impress/ProcedureInfo?action=list&procID=601.

The measurement of the lateral tibia was performed using a calliper after organ extraction. The mouse’s hind limb was bent, and the length measured from the midpoint of the kneecap to the ankle joint^16^. The recorded tibial length, in millimetres (mm), was automatically transferred to the database using a digital or wireless calliper. Data were then uploaded to the IMPC data coordination centre for quality control, processing and analysis prior being released on the IMPC portal.

### Statistical methods

Bespoke methods were developed to assess HR, BW and TL normalization models and are independent of the methods implemented on the IMPC portal.

Data analysis was conducted using *R* (version 4.2.2, R Core Team 2022) with figures and tables produced using *ggplot2* and *ggpubr*. We describe the statistical methods applied to analyse two data groups in this study. ***Real data*** refers to measured data collected from IMPC mice, while ***simulated data*** consists of randomly generated values drawn from a normal distribution in the modelling approach.

#### Real data

Visual methods, as well as formal statistical tests, specifically the Shapiro Wilk test^17^, were applied to test whether the scores of the individual parameters were normally distributed (results not shown). Data were stratified by age and sex and histograms for each parameter were plotted. Reference ranges were calculated based on median, 2.5th percentile, and 97.5th percentile. In addition, the mean, standard deviation, and sample size for each parameter were provided to reflect the data distribution. To explore the linear relationships of HW, BW and TL, the Pearson correlation coefficient (Pearson’s r^18^) was calculated, stratified by sex and age. The strength of these relationships was defined based on user guidelines^19,20^ using the following thresholds: 0.00-0.10 negligible, 0.10-0.39 weak, 0.40-0.69 moderate, 0.70-0.89 strong and 0.90-1.00 very strong correlations. The effects of sex (female vs male) and age (EA vs LA) were compared using identical statistical analyses. In each case, a simple two-tailed t-test was performed, and Cohen’s d effect size was calculated using the *effsize* package. Overinterpretation of statistical significance in large samples is common. Thus, we additionally calculated the effect sizes to better estimate the biological relevance.

#### Simulated data

To compare two independent samples (Group 1 vs. Group 2) in the modelling, we used a Student’s t-test (Student 1908), assuming data drawn from a normal distribution. This test determines whether the two population distributions differ in their parameters, *P_1_* and *P_2_* (**Table 1, Panel A and B**). The following simulation scenarios were tested: Case 1A: *P_1_* and *P_2_*exhibit a linear but non-proportional relationship. Both *P_1_* and *P_2_* are significantly lower in Group 2 compared to Group 1. Case 1B: *P_1_* and *P_2_* exhibit a linear and proportional relationship. Both *P_1_* and *P_2_* are significantly lower in Group 2 compared to Group 1. Case 1C: *P_1_* and *P_2_* are independent and exhibit a non-proportional, random relationship. *P_1_* is significantly lower in Group 2, while *P_2_* is comparable between the groups.

**Table 1:** Simulated data from two independent samples (Group 1 and Group 2) were used in the modelling Cases 1A-1C to determine if their population distributions differ. Panel A: Detailed description of cases and Panel B: The dependence of parameter 2 (*P_2_*) and 1 (*P_1_*).

**Panel A:**
| Simulation Case | Group | Sample Size | Parameter 1 | Parameter 2 | Error term $\epsilon$ |
| --- | --- | --- | --- | --- | --- |
| Case 1A | Group 1 | n=10 | $\mathcal{N}(35,3)$ | $30+(1.5*\text{Parameter 1}) + \epsilon$ | $\mathcal{N}(0,3)$ |
| | Group 2 | n=10 | $\mathcal{N}(30,3)$ | $30+(1.5*\text{Parameter 1}) + \epsilon$ | $\mathcal{N}(0,3)$ |
| Case 1B | Group 1 | n=10 | $\mathcal{N}(35,3)$ | $1.5*\text{Parameter 1} + \epsilon$ | $\mathcal{N}(0,3)$ |
| | Group 2 | n=10 | $\mathcal{N}(30,3)$ | $1.5*\text{Parameter 1} + \epsilon$ | $\mathcal{N}(0,3)$ |
| Case 1C | Group 1 | n=10 | $\mathcal{N}(35,3)$ | $\mathcal{N}(80,5)$ | |
| | Group 2 | n=10 | $\mathcal{N}(30,3)$ | $\mathcal{N}(80,5)$ | |

**Panel B:**
| Case | How $P_2$ is determined |
| --- | --- |
| **1A** | Linear but not proportional to $P_1$ |
| **1B** | Linear and proportional to $P_1$ |
| **1C** | Random with no connection to $P_1$ |

### Linear model

A linear model is a statistical approach used to describe the relationship between a dependent variable (*Y*) and one or more independent variables (*X*)^21^. It assumes that this relationship can be expressed as a linear equation. Here, a linear model was applied using the *lm()* function to perform a simple linear regression between HW and BW.

### Allometric scaling

Allometric scaling follows a power-law relationship rather than a simple linear trend^22^. The allometric model is typically expressed as:

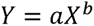

where *a* is the scaling constant and *b* is the scaling exponent.

As this equation is nonlinear, we applied a log-log transformation to linearize it, enabling the use of linear regression to estimate the allometric exponent *b*:

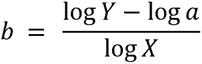

In this study, the coefficient *b* indicates how HW scales with BW or TL. Allometric scaling relationships between HW and BW, as well as between HW and TL, were generated.

All models and the *R*-code are openly available in our GitHub repository: https://github.com/ExperimentalGenetics/AllometricScaling.

### Data availability

All data used in this study are publicly available from the IMPC (https://www.mousephenotype.org/data/release).

## Results

Heart weight (HW) data collected by the IMPC contributing centres (data release, DR, 21.0) are available from 25,354 wildtype mice. In the EA population, 27 mice were excluded from analysis due to age inconsistency. The remaining 25,327 mice were stratified as presented in **Table 2** and summarized below. All mice are from a C57BL/6N inbred substrain. Most mice (86.3% or 21,846) were tested at a median age of 16 weeks (designated as “early adult” or EA), while the remaining 13.7% (3,481 of mice were tested at a median age of 60 weeks (designated as “late adult” or LA). Sex is evenly distributed at both the EA and LA time points. Body weight (BW) data are available for all mice whereas tibia length (TL), a non-mandatory IMPC parameter, is available for 7,775 (35.6% of 21,846) EA and 1,431 (41.1% of 3,481) LA mice. The total number of parameters reported varies slightly between mice and can be accessed in each table.

**Table 2:** Heart weight data were available from a total of 25,327 mice, stratified by sex and age at testing (EA = median of 16 weeks of age; LA = median of 60 weeks of age). Body weight was recorded for all mice; however, tibia length measurements were not uniformly available across the dataset.

| Age | EA | EA | LA | LA | Total |
| --- | --- | --- | --- | --- | --- |
| Sex | female | male | female | male |  |
| *Heart Weight [mg]* | N = 11016 | N = 10830 | N = 1782 | N = 1699 | N = 25327 |
| *Tibia Length [mm]* | N = 3890 | N = 3885 | N = 724 | N = 707 | N = 9206 |

### Evaluating data distribution: metrics and significance testing

The distribution of data was assessed using histograms for HW, BW and TL stratified by sex and age (**Figure 1, Panel a-c**). Calculated ranges using mean ± standard deviation (SD), as well as median and 95% reference ranges (2.5th and 97.5th percentiles) were additionally provided for consistency (**Figure 1, Panel d-e**). The visual representation of value frequencies was useful in illustrating both conformity to and deviations from a normal distribution for each parameter. Visual inspection of the histograms showed that the data appeared practically normally distributed in both EA and LA groups for male and female mice.

**Figure 1:**
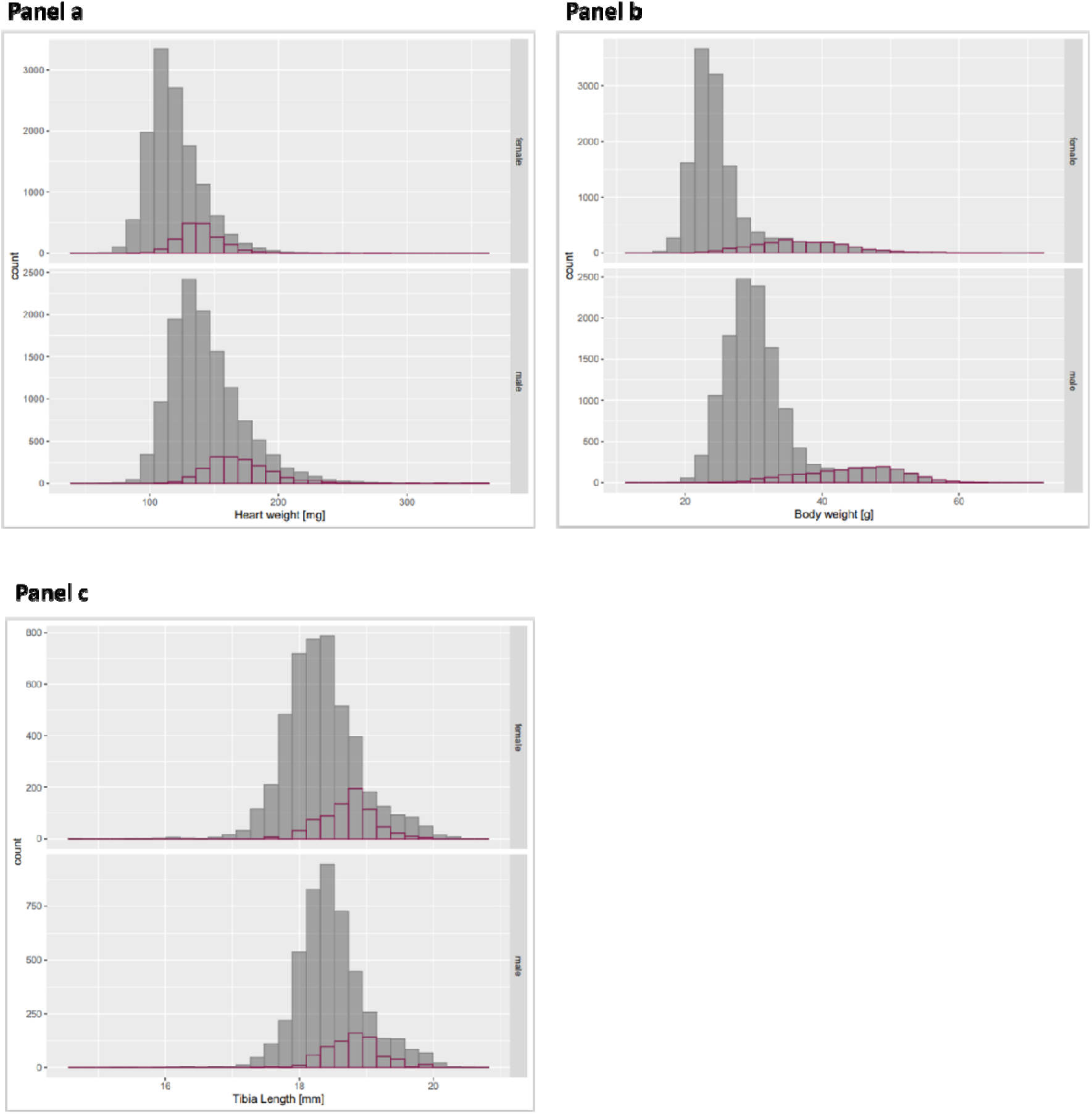
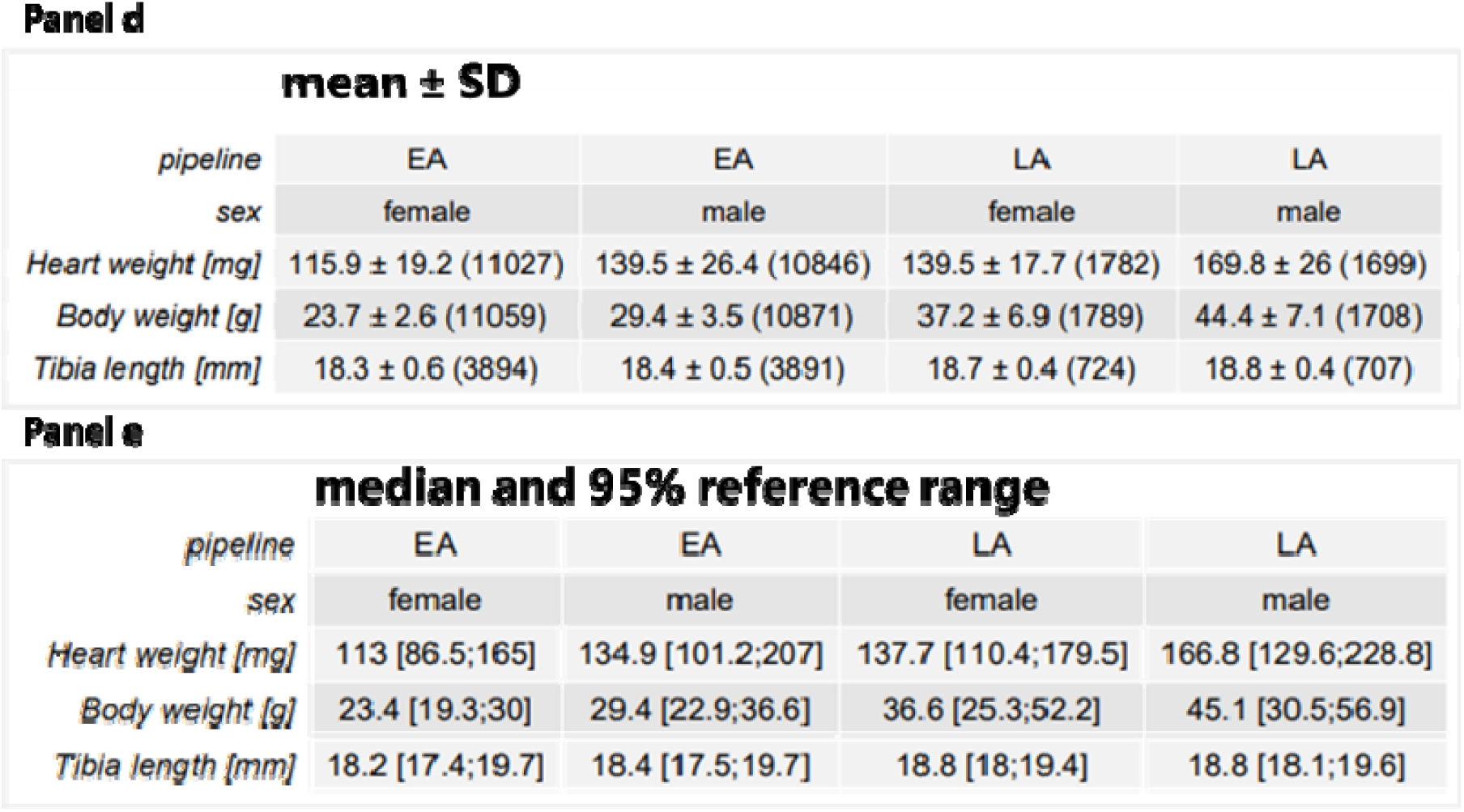
Histograms showing the distribution of heart weight [mg], body weight [g] and tibia length [mm] data along with tables depicting calculated ranges (mean ± SD and median with 95% reference ranges) for EA (grey) and LA (purple) mice, stratified by sex. These calculations are based on data from eight contributing IMPC centres. Panel A depicts heart weight, Panel B body weight and Panel C tibia lengths for EA (**grey**) and LA (**red**) mice, stratified by sex. Panel D shows mean ± SD and Panel E median with 95% reference ranges for EA and LA mice, stratified by sex.

**Figure 1** shows that EA male and female data showed similar distributions by visual inspection with males having higher values for HW, BW and similar TL compared to females. The same pattern was observed for the LA population. When comparing the LA to EA data sex-specifically, both LA females and LA males showed higher HW, BW and TL values than the corresponding EA populations (**Figure 1**).

To test the hypothesis that there is a difference between sexes, a simple two-tailed t-test was performed independently for each age group (EA vs LA), and Cohen’s d was calculated as an effect size measure. In both the EA and LA populations, for HW, BW and TL, p-values reached significance (≤.001). Moreover, for all parameters, the corresponding Coheńs d value revealed large effect sizes between sexes for HW (1.03-1.37) and BW (1.02-1.87), whereas TL showed minor (0.29) to negligible (0.17) effect sizes in EA and LA mice, respectively (**Table 3, Panel A**).

**Table 3:**
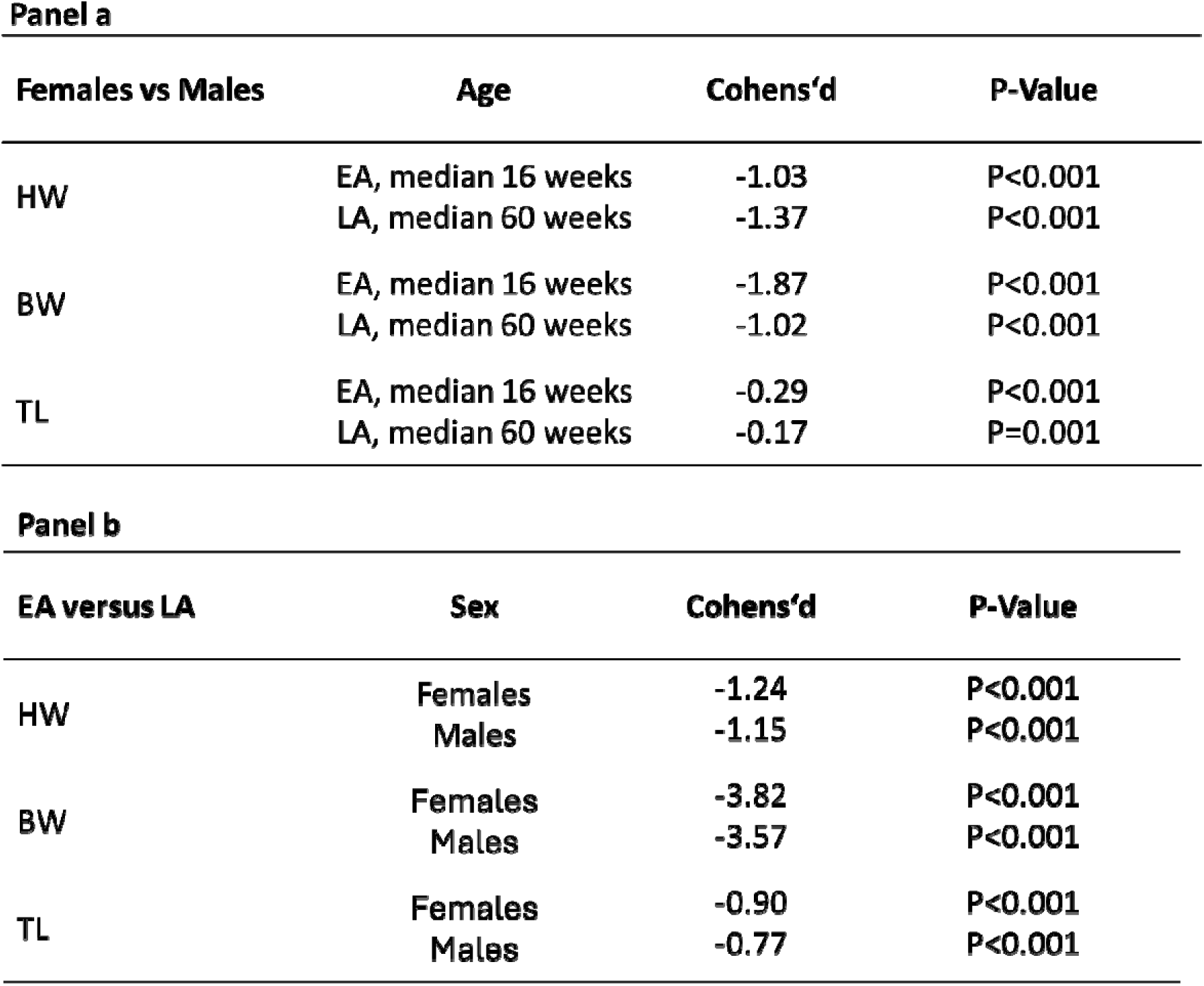
Sex-specific differences in heart weight (HW), body weight (BW), and tibia length (TL) were evaluated in early adult (EA) and late adult (LA) groups. Two-tailed t-tests showed significant differences (≤.001) for all parameters, with large effect sizes for HW and BW and minor to negligible effects for TL based on Cohen’s d values (Panel a). Age-related comparisons also revealed significant differences (≤.001) with medium to large effect sizes for HW, BW, and TL, stratified by sex (Panel b).

**Panel a**
| **Females vs Males** | **Age** | **Cohens'd** | **P-Value** |
| --- | --- | --- | --- |
| HW | EA, median 16 weeks | -1.03 | P<0.001 |
|  | LA, median 60 weeks | -1.37 | P<0.001 |
| BW | EA, median 16 weeks | -1.87 | P<0.001 |
|  | LA, median 60 weeks | -1.02 | P<0.001 |
| TL | EA, median 16 weeks | -0.29 | P<0.001 |
|  | LA, median 60 weeks | -0.17 | P=0.001 |

**Panel b**
| **EA versus LA** | **Sex** | **Cohens'd** | **P-Value** |
| --- | --- | --- | --- |
| HW | Females | -1.24 | P<0.001 |
|  | Males | -1.15 | P<0.001 |
| BW | Females | -3.82 | P<0.001 |
|  | Males | -3.57 | P<0.001 |
| TL | Females | -0.90 | P<0.001 |
|  | Males | -0.77 | P<0.001 |

Two different age groups, i.e. median age of 16-weeks (minimum 15 and maximum 20 weeks) old EA mice and median age of 60 weeks (minimum 55 and maximum 81 weeks) old LA mice, made it possible to explore the effect of age on HW, BW and TL, stratified by sex. P-values ≤.001 were reached for all parameters, indicating high statistical significance. The corresponding Cohen’s d effect size values revealed medium to large, standardized effect sizes in HW, BW and TL (**Table 3, Panel B**).

### Examining correlations: heart weight in relation to body weight and tibia length

To explore the linear relationship between HW and either BW or TL, we calculated the Pearson correlation coefficient *r*, stratified by sex and age. Female data are placed directly next to male data for ease of visualization.

**Figure 2** shows distinct distribution clusters with regression lines for EA and LA groups stratified by sex. Negligible to weak standardized Pearson correlations were obtained for both comparisons in the EA population with HW (mg) and BW (g) showing r=0.21 for females and r=0.21 for males (**Figure 2**, **Panel A & B**). In the LA population, the correlations were r=0.26 for females and r=-0.069 for males (**Figure 2, Panel A & B**), indicating a weak or negligible correlation between HW and BW; the HW (mg) to TL (mm) ratio reached in the EA population r=0.058 for females and r=0.014 for males (**Figure 2**, **Panel C & D**), whereas in the LA population it was r=0.055 for females and r=-0.12 for males (**Figure 2, Panel C & D**). Comparing EA and LA populations, the relationship flattens over time, indicating a non-linear trend across the lifespan from EA (median 16 weeks) to LA (median 60 weeks) in healthy C57BL/6N substrain mice (**Figure 2, Panel A-D**).

**Figure 2:**
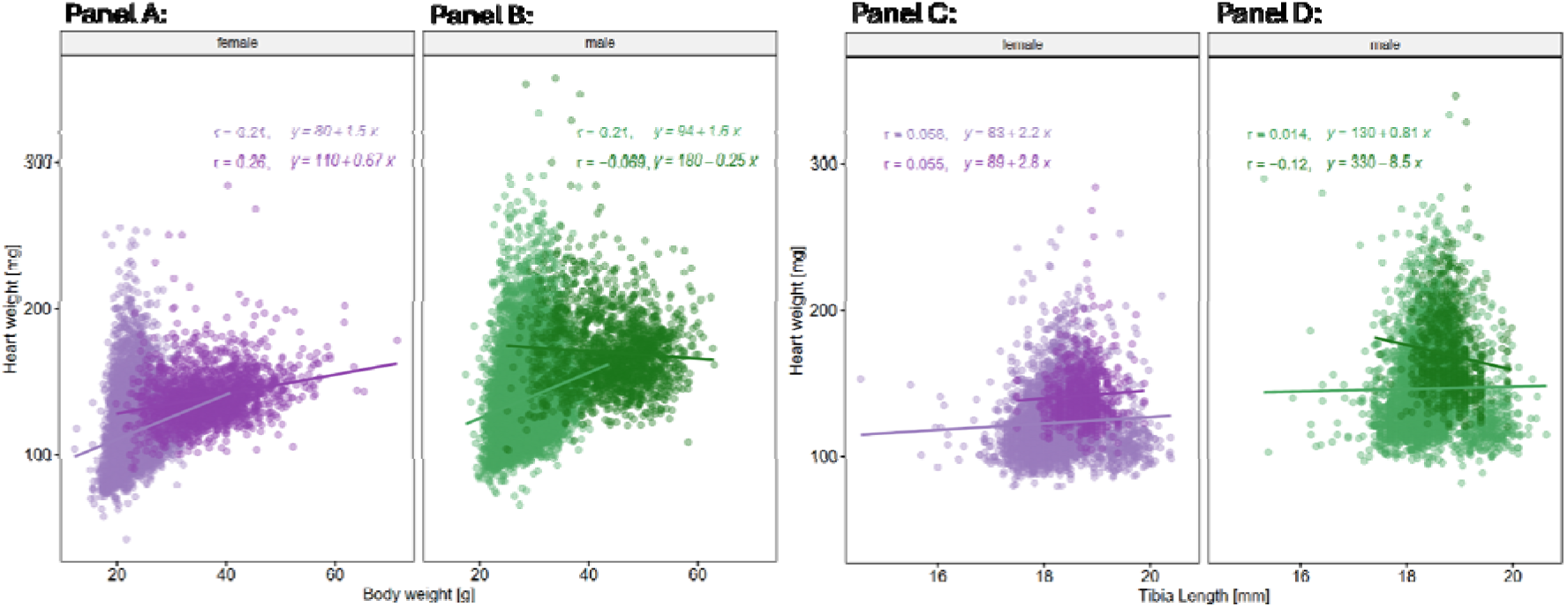
The linear models in the plot are represented by the regression equations displayed on each Panel. These equations describe the relationship between the independent variable (body weight [g] or tibia length [mm]) and the dependent variable (heart weight [mg]) for both female and male groups. The data are stratified by sex (female vs. male) and age group (early adult [EA] vs. late adult [LA]). Colour codes are as follows: **light purple** for early adult (EA) female and **light green** for early adult (EA) male mice, **dark purple** for late adult (LA) females and **dark green** for late adult (LA) males. Panel A: HW with BW in EA (**light purple, r=0.21**) and LA (**dark purple, r=0.26**) female populations; Panel B: HW with BW in EA (**light green, r=0.21**) and LA (**dark green, r=-0.069**) male populations. Panel C: HW with TL in EA (**light purple, r=0.058**) and LA (**dark purple, r=0.055**) for females; Panel D: HW with TL in EA (**light green, r=0.014**) and LA (**dark green, r=-0.12**) for males.

### Unveiling biases: modelling the limitations of ratio-based analyses in non-proportional relationships

Following on from the non-proportional relationship shown here, we aim to demonstrate through artificial modelling that using ratios to compare groups is mathematically inconsistent.

We conducted a simulation study with artificial data to demonstrate that parameter ratios, when used to detect differences between groups, can lead to misleading conclusions and should only be applied under specific conditions. Simulated datasets were generated for two groups, group 1 and group 2, each consisting of 10 samples (**Table 1, Panel A**). The values for two parameters, parameter 1 and parameter 2, were used to reflect modelling scenarios across different cases (**Table 1, Panel B**). Parameter 1 (*P_1_*) was drawn from a normal distribution. Depending on the case, parameter 2 (*P_2_*) was either derived from *P_1_* through a defined relationship (non-linear or proportional) or independently sampled from a normal distribution with predefined mean and standard deviation.

Case 1A: *P_2_* is related to *P_1_* but in a non-proportional way. This suggests a non-linear (or non-constant scaling) relationship such as: *P_2_*= *f (P_1_) +* L where *f (P_1_)* is a non-proportional function (*P_2_*≠c⋅*P_1_*), and L is an error term.

Case 1B: *P_1_*is approximately proportional to *P_2_* with *c* the constant scaling factor, expressed as *P_2_*= *c*⋅*P_1_ +* L, where L is an error term.

Case 1C: *P_2_*is independently drawn from a normal distribution where *P_2_* does not depend on *P_1_* and is randomly sampled with μ*_2_* and σ ^2^ independent of *P_1_*: *P_2_*= *N (*μ*_2_,* σ ^2^).

A t-test was used to compare differences between the two groups, with significance evaluated at a threshold of ≤.05. **Table 1** provides model details.

The results in **Figure 3** show that in Case 1A applying the ratio of *P_1_*to *P_2_* retains a statistically significant difference between group 1 and group 2 (≤.05) but significantly reverses the direction of the effect compared to the raw data. This indicates that the ratio causes distortion under these conditions. In contrast, in Case 1B *P_1_* and *P_2_* yet show a linear and proportional relationship. Using the ratio in this case eliminates group differences and yields results consistent with those obtained from the linear model with *P_1_* as a covariate. This shows that the ratio only applies in this specific context, where the scaling relationship between the two parameters is appropriately controlled. In Case 1C both parameters are simulated independently from each other and show a random non-proportional relationship. Calculating the ratio of *P_2_*to *P_1_* gives a false significant difference for *P_2_*, although there is no real difference between groups. This discrepancy observed in Case 1C arises because the regression line does not pass through the origin, thereby violating a fundamental assumption required for the valid use of ratios.

**Figure 3:**
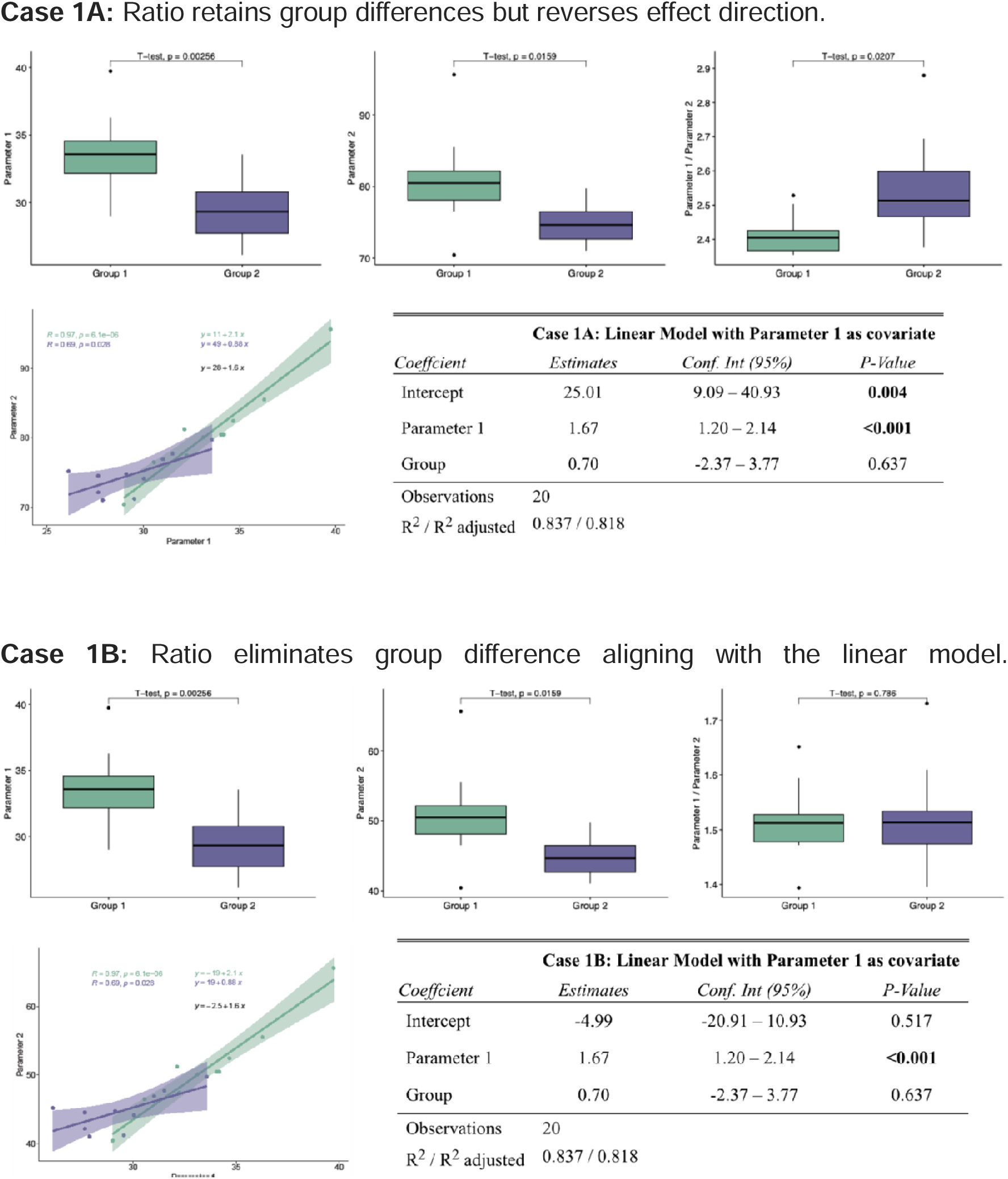
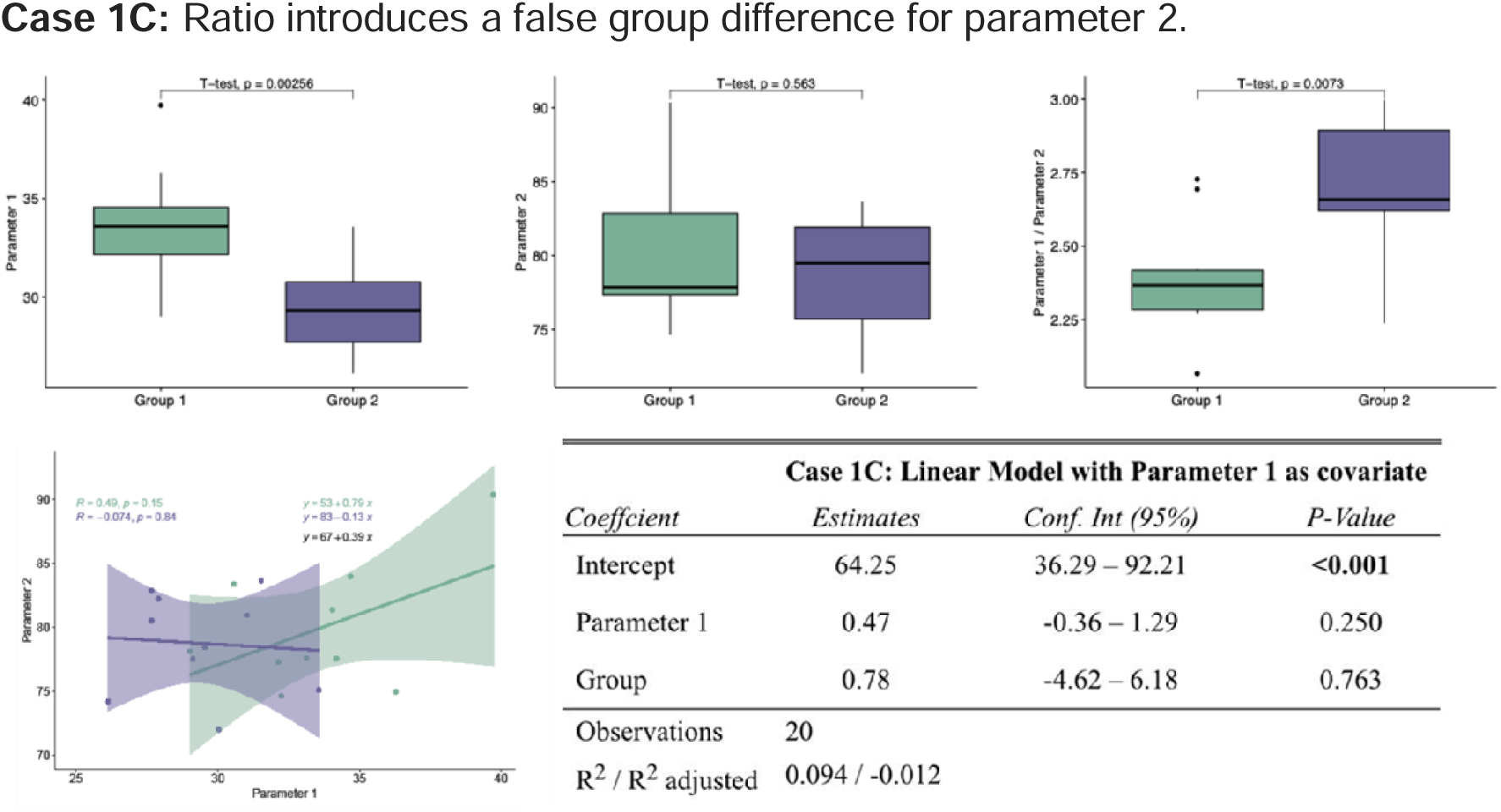
Modelling demonstrates the mathematical inconsistency of non-proportional relationships when comparing groups. Simulated datasets for two groups (n=10 per group) were generated with values for two parameters randomly drawn from a normal distribution. A t-test (≤0.05) was applied to compare groups with parameter 1 and 2, highlighting that ratio-based analyses can lead to incorrect conclusions and are only valid under specific conditions (Case 1B). Colour codes are as follows: **green** for group 1 and **purple** for group 2. **Case 1A:** Ratio retains group differences but reverses effect direction. **Case 1B:** Ratio eliminates group difference aligning with the linear model. Because of the additional error term □∼□(*0,3)*, the regression lines are not perfectly proportional.

Applying this simulated modelling approach to our data set, the conclusions remain identical. This implies that regardless of the body size parameter considered - whether TL or BW - the fundamental methodological challenges and scaling issues persist.

Simple ratio-based normalization methods often do not capture the underlying biological relationships. In comparison, linear and allometric models offer a more robust and precise analytical framework.

### Beyond ratios: a linear modelling approach for more robust biological scaling analysis

A linear model (LM) provides a simplified but powerful way to analyse and predict relationships between variables in both statistics and scientific research. Here, an LM assumes a direct, additive relationship between HW and BW, allowing for the estimation of changes in HW as a function of BW. This approach accounts for variability by including an intercept and a slope term, making it a more flexible alternative to simple ratio-based normalization. The formula for the linear model is *y = a + bx*, where *y* represents heart weight (mg), and *x* denotes either body weight (g) or tibia length (mm). The intercept *a* is the extrapolated heart weight when *x=0*, although this may not be biologically meaningful. The slope *b* represents the predicted change in heart weight (mg) per unit increase in *x* (g or mm). **Figure 2** shows the LM with regression lines for the relationship between HW and BW in early adult (EA) and late adult (LA) females (Panel A), and in EA and LA males (Panel B). Panels C and D show the relationship between HW and TL for EA and LA females and males. The plots differentiate between males and females, showing how heart weight relates to body weight and tibia length in each group. Furthermore, differences in slope (*b*) suggest that the relationship between parameters differs between sexes. Most equations (Panel A-C) have positive slopes, meaning HW increases as BW or TL increases. However, in Panel D, the male TL has a negative slope (-8.5), indicating an inverse relationship in the LA male population.

In summary, the LM is visualized in **Figure 2** through regression equations and their corresponding trend lines, highlighting how HW depends on BW or TL in both males and females. The variation in slopes suggests sex-specific linear relationships.

### From linear models to allometric scaling: a scientific transition

Initially, we modelled the relationship between HW and BW using a linear regression model, assuming a constant rate of change. However, organ weights often do not scale linearly with body weight.

To account for nonlinear growth patterns, we adopted an allometric scaling model, which is commonly used in biological systems to describe how physiological traits change with body size^23^. This model follows the power-law equation: *y = ax^b^*, where the exponent *b* determines the scaling pattern. In this equation, *y* represents HW, *x* denotes BW, *a* and *b* represent the scaling coefficient and the allometric exponent respectively, which together define the scaling relationship.

The allometric exponent *b* provides insights into the growth rate of HW relative to BW. When *b=1*, isometric scaling occurs, meaning that HW increases proportionally to BW. A value of *b<1* suggests negative allometry, where HW increases at a slower rate than BW, while *b>1* indicates positive allometry, where HW increases at a faster rate than BW.

**Figure 4** shows that our findings (b=0.38 for females, b=0.39 for males) indicate negative allometry, meaning HW increases at a slower rate than BW in a healthy wildtype population. This shift from a linear to an allometric approach provides a more accurate and biologically meaningful representation of cardiovascular scaling.

**Figure 4:**
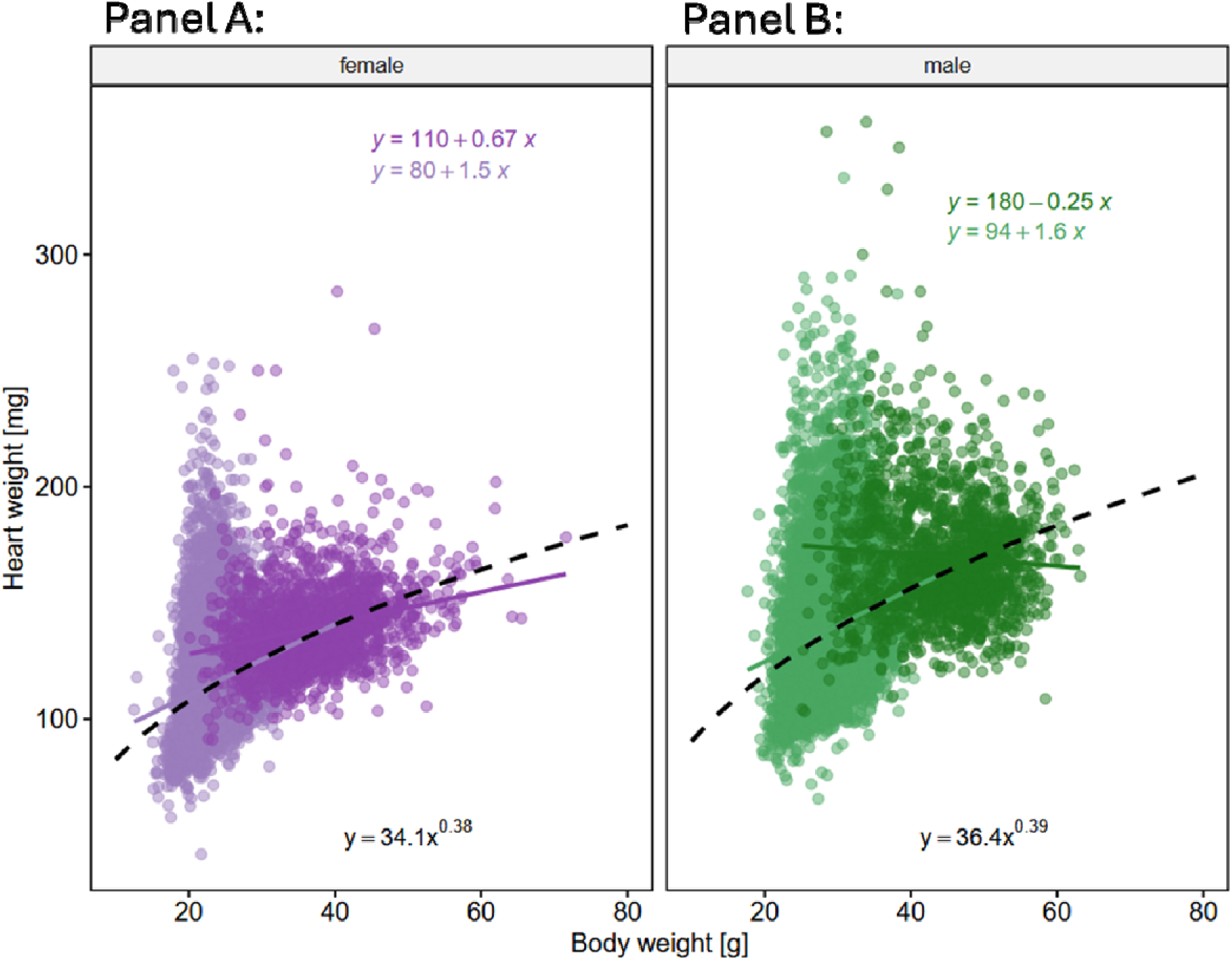
Allometric model for body weight [g] and heart weight [mg] represented by the regression equations in blue. Data are stratified by sex and age. Female mice in early adult (EA) population are represented in **light pink**, while late adult (LA) females are shown in **dark pink** (Panel A). Male mice in the EA group are depicted in **light green** and LA males in **dark green** (Panel B).

### Scientific justification for using allometry

The observed allometric exponents in our study, b=0.38 for females and b=0.39 for males, indicate negative allometry, which is consistent with previous findings in cardiovascular physiology^23^. For example, studies such as Prothero^24^ and Calder^25^ have shown that mammalian heart weight typically scales with body weight with exponents below 1, reflecting negative allometry as animals increase in size. Similar sublinear scaling exponents were reported for cardiac tissue relative to body mass and metabolic rate^26^. This suggests that, according to allometric scaling laws, as body size increases, the heart does not grow at the same rate but rather adapts to maintain efficiency in circulatory function. Such relationships align with broader biological scaling principles, such as those described by Kleiber and Cole^27^, which explains how metabolic rates scale sublinearly with body mass.

## Discussion

The results presented here, supported by the extensive dataset provided by the International Mouse Phenotyping Consortium, emphasize the power of large-scale, standardized data to address methodological questions in quantitative biology. This study highlights the limitations of ratio-based analyses in allometric datasets. It underscores the importance of using linear models with covariate adjustment or even allometric scaling models to generate robust and interpretable results, depending on the data structure.

### Biological context: allometry and ratios

In normal physiological conditions, organ weights, such as heart weight (HW), do not always exhibit strong linear and proportional relationships with metrics like body weight (BW) or tibia length (TL). This study shows that in wildtype C57BL/6N mice, correlations between HW and BW, and HW and TL, are weak to negligible across sex and age groups. Such weak non-proportional relationships inherently undermine the validity of ratio-based normalization (e.g., HW/BW or HW/TL) in detecting group differences, as these ratios assume proportional scaling between variables, which is not supported by the data.

The observed variability in HW relative to BW and TL also reflects the biological reality that organ development and body size are influenced by different environmental and physiological factors. This underscores the need for analytical approaches that account for these complexities without introducing artifacts^28^.

### Modelling study: pitfalls of ratio-based normalization

Through simulated data, we demonstrated how using ratios under weak linear or proportional relationships can lead to mathematically inconsistent conclusions when comparing groups. In Case 1A, applying a ratio preserved statistically significant group differences but reversed the direction of the effect, leading to misleading conclusions. Conversely, in Case 1B, where specific assumptions (e.g., a regression line passing through origin) were met, the ratio produced accurate results comparable to those from linear models. However, Case 1C demonstrated that ratios can artificially create significant differences between groups even when none exist, a direct consequence of invalid scaling assumptions.

These findings reinforce that ratio-based comparisons between groups are only appropriate in rare, specific cases where the relationship between variables is both linear and proportional, that is when the regression line passes through the origin. Otherwise, ratios risk distorting results, as demonstrated in this study.

### Rethinking ratios: considerations for and limitations of large-scale phenotyping

The IMPC dataset, encompassing over 25,000 wildtype mice, offers unmatched statistical power and granularity for systematically investigating allometric relationships. Utilizing this extensive resource, we demonstrate that linear or power-law based models outperform ratio-based approaches in analysing allometric data. Unlike ratios, these models maintain the integrity of the underlying biological relationships, adjust for confounding variables, and provide statistically robust and biologically meaningful conclusions.

No study is without limitations; here, we cannot exclude that the large-scale multi- centre setup introduces centre effects and parameter variations that may influence the results. Looking ahead, including mutant mouse lines and distinguishing whole-body weight from body composition will help assess non-allometric scaling relevant for compensatory or pathological hypertrophy in the heart. We modelled three scenarios here, but they do not cover all possible real-world situations, so results may vary depending on the biological context. In consequence, the key takeaway is that visual inspection together with preliminary data review are essential for the selection of appropriate models to draw meaningful conclusions.

### Guiding future research: key recommendations based on findings

- Ratio-based comparisons between groups should be avoided unless the relationship between parameters is strictly linear and proportional, with a regression line passing through the origin.
- Including a scatterplot representation of the data could provide a clearer visualization of the relationship, as it offers a meaningful way to explore the data structure before applying statistical models.
- Linear and power-law models with covariate adjustments should be the preferred methods for analysing allometric relationships, as they control confounders and accommodate certain non-proportionality.
- The use of large-scale datasets, such as those generated by the IMPC, should be encouraged to validate analytical approaches and ensure reproducibility across diverse biological contexts.

### Conclusion: Why the shift matters

Transitioning from a linear model to an allometric scaling model provides a more biologically meaningful interpretation of how heart weight varies with body weight. This approach better captures nonlinear growth dynamics, allowing a more accurate representation of physiological constraints across different body sizes.

Moreover, this study provides compelling evidence against the indiscriminate use of ratio-based comparisons in allometric analyses and emphasizes the critical importance of appropriate statistical methods to ensure valid and interpretable results. The IMPC’s extensive phenotyping data serve as a powerful resource for refining analytical practices in quantitative biology, ultimately advancing our understanding of complex biological systems.

## Author contributions

Mouse Production: Susan Marschall

Phenotyping & Data Acquisition: Patricia da Silva-Buttkus; Helmut Fuchs

Data Analysis: Manuela A Oestereicher; Elida Schneltzer

Manuscript Design & Preparation: Manuela A Oestereicher; Nadine Spielmann

Design & Study Supervision: Manuela A Oestereicher; Elida Schneltzer; Martin Hrabě de Angelis; Nadine Spielmann

PI on grant: Martin Hrabě de Angelis

Review the manuscript: Elida Schneltzer; Valérie Gailus-Durner; Martin Hrabě de Angelis (MHdA)

## IMPC consortium

Heaney Jason D.^1^, White Jacqueline K.^2^, Herault Yann^3^, Tamura Masaru^4^, Lloyd K.C. Kent^5^, Seong Je Kyung^6^, Nutter Lauryl M. J.^7^

^1^Department of Molecular and Human Genetics, Baylor College of Medicine, One Baylor Plaza, Houston, Texas, 77030, United States of America

^2^Center for Biometric Analysis, The Jackson Laboratory, 600 Main Street, Bar Harbor, ME USA 04609

^3^Université de Strasbourg, CNRS, INSERM, Institute de la Clinique de la Souris, PHENOMIN, 1 rue Laurent Fries, 67404 ILLKIRCH, France

^4^Experimental Animal Division, RIKEN BioResource Research Center,3-1-1 Koyadai, Tsukuba, Ibaraki 305-0074, Japan

^5^Mouse Biology Program, 2795 Second Street Suite 400, University of California, Davis, 50 California 95618, USA

^6^Korea Mouse Phenotyping Consortium (KMPC) and BK21 Program for Veterinary Science, Research Institute for Veterinary Science, College of Veterinary Medicine, Seoul National University, Seoul, South Korea

^7^The Hospital for Sick Children, Toronto, ON M5G 1X5, Canada

## Acknowledgements

This work would not have been possible without the support of the IMPC phenotyping centres. Hands-on mouse IMPC phenotyping has been carried out by many laboratory staff with superb experimental skills and unsurpassed dedication. The authors thank these individuals for their contribution to the IMPC data used herein.

